# The Role of Synchronization of Neural Modules in Pattern Processing in the Visual System

**DOI:** 10.64898/2025.12.29.696937

**Authors:** O.V. Levashov, V.F. Safiulina

## Abstract

Key operations in modeling visual processing in living systems are those that process a pattern as a holistic object, for example, in pattern recognition to ensure invariance to translation and rotation. In this article, we describe a neural module we synthesized for moving a pattern along a cortical layer without losing its shape. The module has the form of a three-layer neural network with two types of inhibitory neurons, one of which is a universal “prohibition” logic element. Modeling showed that such operations can be implemented using the proposed neural module by additionally synchronizing its operation using endogenous pacemakers with a gamma rhythm frequency. The mechanism of such synchronization is similar to the clock sweep mechanism in electronics. We believe we have clearly demonstrated the possible role of gamma-based brain activity as a mechanism for synchronizing computations in living neural networks for the first time. This opens up new possibilities for constructing neuromorphic architectures for real-time visual recognition.

## Introduction

One of the key conditions for reliable visual perception and recognition in living systems is the maintenance of an unchanging form of “mental patterns” at the level of cortical neural networks. If a pattern is distorted during a short period of eye fixation, the recognition task will be greatly hampered. This problem is often overlooked when modeling recognition in living visual systems. However, there are a number of facts that indicate that the problem of pattern blurring at the level of cortical neurons does exist.

Indeed, neurons in the cerebral cortex are interconnected by a huge number of connections, which is clearly visible even in morphological sections of the cortex. The fact that cortical neurons are “multiconnected” is also confirmed by changes in the nature of their electrical activity during early development. For example, giant potentials arise in rats on the first or second day after birth, but their amplitude soon normalizes (Ben-Ari et al., 1989). This effect is explained by rapidly emerging connections between neurons and the synchronization of their spiking activity (Safiulina et al., 2004). The instability of the pattern in the visual cortex may also be related to the random nature of the action potential generation process. It is known that each neuron in the resting state exhibits spontaneous activity, and its input and output characteristics are, to a certain extent, random variables and can change over time (Amir et al., 1993; Bair and Koch, 1996; Butts et al., 2007; Liu et al., 2001; Nolte et al., 2019; Reich et al., 1997; Tolhurst et al., 1983).

An important point for us is the idea that neurons in a working neural network, although connected, from a mathematical perspective remain discrete sets. Therefore, we hypothesized that synchronization of the neurons involved in processing such a pattern of neural activity is necessary to perform the needed manipulations.

Therefore, the goal of our study was to determine:

1. How stable is the pattern during processing in living neural networks in the 300-500 ms interval, i.e., during the eye fixation period between two saccades (Yarbus, 1965). If it blurs or otherwise changes shape, what might be the mechanism for its “freezing” (fixation)?
2. Is it possible to synthesize modules from excitatory and inhibitory model neurons that could perform image processing operations in vision?
3. How can we synchronize the processing of a pattern represented in the cortex as a set of activated neurons, that is, essentially not a complete pattern due to the discrete nature of the neurons themselves?

We found no descriptions of similar working neural networks in the literature. Therefore, in this article, we describe the neural module we developed, a three-layer neural network with two types of inhibitory neurons, one of which is a universal “prohibition” logic element. Furthermore, we propose a mechanism for synchronizing the operation of these neural modules based on endogenous pacemakers, which, similar to electronic devices, generate clock frequencies for processing input stimuli.

### Modeling the preservation of input pattern shapes in visual recognition

The simulation was performed on a 50×50 raster, where each cell corresponded to one neuron in the network. The full codes are available in the repository (https://github.com/victoriasafiulina-design/ModelV). The connections between all neurons were stochastically defined, with connection radii of 1, 2, and 3 cells and a probability of 0.3. The proportion of inhibitory neurons was 20%.

In the first experiment, all neurons were excitatory.

In the second experiment, 20% inhibitory neurons were added.

In the third experiment, a special 3-layer neural network was simulated, including, in addition to standard inhibitory neurons, inhibitory neurons of the PROHIBITION type. Furthermore, in this case, the synchronous action of an endogenous pacemaker on the entire lower layer of inhibitory neurons was simulated.

The fourth experiment used the same three-layer neural network, but added temporal synchronization via a single bus with endogenous pacemakers. When the right pacemaker was activated at a gamma frequency (100 Hz), the stimulus pattern shifted as a whole to the right, while when the left pacemaker was activated, it shifted to the left.

### Simulation results

In the first experiment, it was shown that the stimulus pattern quickly lost its shape and became blurred (Fig. 1).

**Fig. 1.**
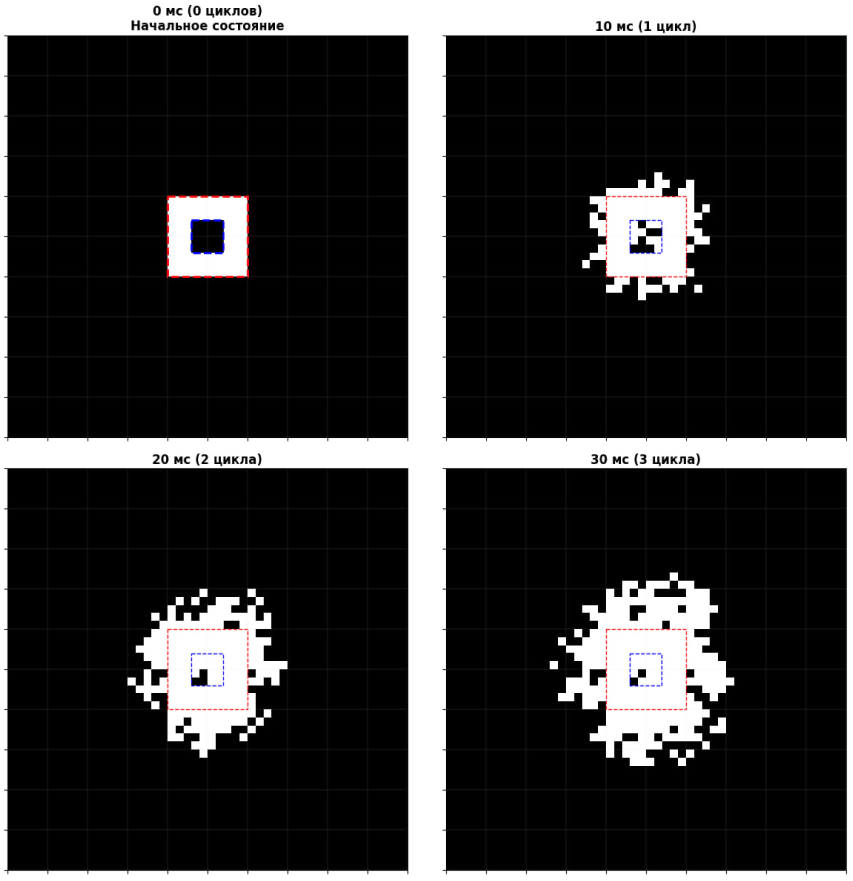
Computer simulation of the “fate” of the original pattern (left, first row) in a field of 50×50 neurons 10, 20, and 30 ms after the initial pattern was applied to this field. All neurons were excitatory. As can be seen, the original pattern becomes blurred after just 20 ms.

However, it is known that the cortex contains inhibitory neurons that could prevent pattern blurring during processing. According to various estimates, such inhibitory interneurons account for approximately 10-20% of the cortex, with many different types being distinguished. For example, the hippocampus contains up to 29 types of inhibitory interneurons (Kasyanov et al., 2004; Kessaris and Denaxa, 2023; Tzilivaki et al., 2023). This diversity of interneurons suggests that they play an important role in signal transmission and processing in the visual brain. Therefore, in the next step of modeling, we introduced inhibitory interneurons into our test neural network (Fig. 2).

**Fig. 2.**
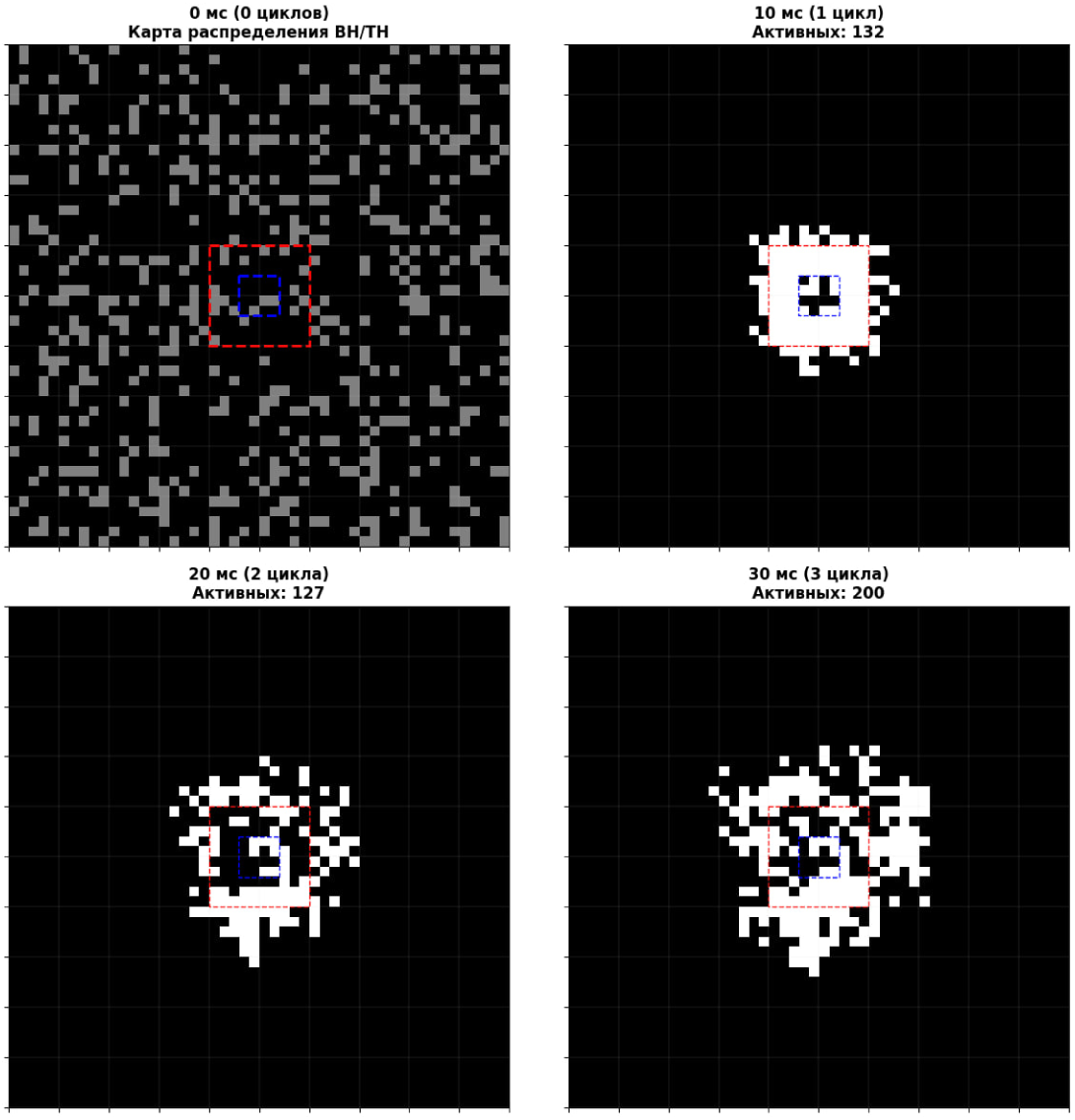
Pattern blurring after introducing randomly placed inhibitory neurons (20% of the total number of neurons). The first image shows the distribution of excitatory (black cells) and inhibitory neurons (gray cells). It is clear that after 30 ms, the original pattern has lost its shape.

In this case, it is clear that despite the presence of randomly placed inhibitory neurons, the pattern has become blurred, not as widely, but blurring precisely at its boundaries, losing both the outer and inner boundaries.

### Modeling a neural module for pattern capture during visual recognition

To prevent blurring of the input neural activity pattern, we constructed a neural network (neural module) consisting of two types of excitatory and inhibitory neurons (Fig. 3):

1. Regular inhibitory neurons, which act on excitatory neurons.
2. “Prohibition” neurons, which interact only with inhibitory neurons of the first type, locking them (disabling) at certain points in time during pattern processing.

**Fig. 3.**
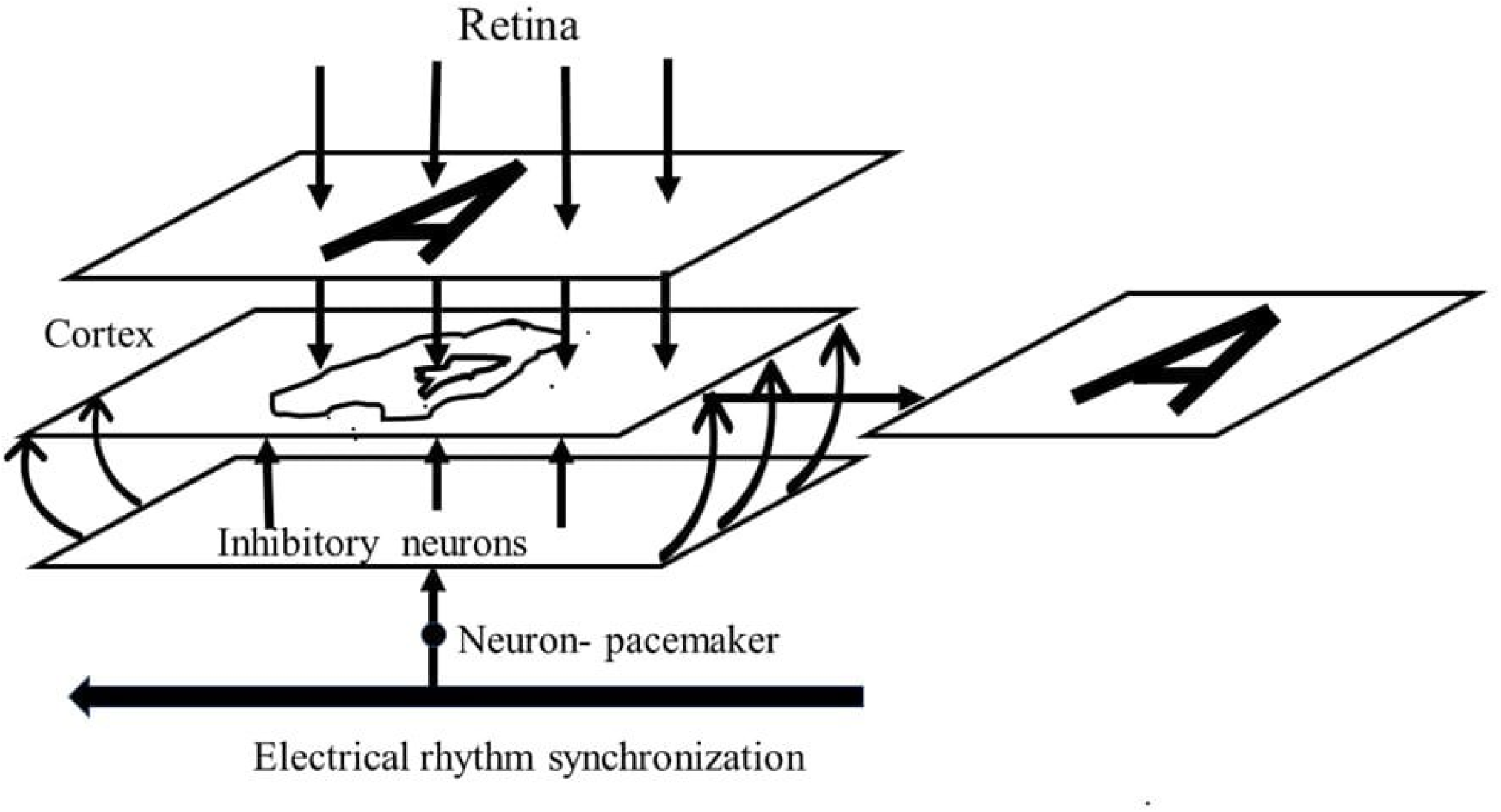
General structure of the neural model for “fixing” a visual pattern during cortical processing. The input image (in the figure, shape A on the retina) is projected onto the “cortical neural layer” (the second plane with a blurred shape). This layer simulates cortical neurons, which are interconnected by a large number of excitatory connections. A layer of inhibitory interneurons (shown further down in the figure) sends inhibitory signals to the upper layer, where the processed pattern resides, at each scan cycle. It also inhibits neurons in this layer at the edges of the pattern, preventing the pattern from blurring (the activity of excited neurons is not transmitted to inactive neurons adjacent to the pattern). At the same time, excited neurons from the active pattern in the second layer have special connections with the lower layer of inhibitory neurons—inhibitory connections. Thus, they disable only those inhibitory interneurons located directly beneath the pattern. As a result, the pattern remains active and does not blur.

All these model neurons belong to the “continuous logic” type of neurons first proposed by N.V. Pozin and his colleagues in Institute of Control Sciences in Moscow (Pozin, 1971; Pozin et al., 1978). Subsequent modeling clearly demonstrated that neurons of these three types, like a construction set, can be used to assemble virtually any neural module implementing any information processing operation (Levashov, 2021).

The presence of different types of inhibitory neurons in the cortex, including “prohibition” neurons, is supported by physiological data. For example, in a study (Pfeffer et al., 2013), different types of interactions were discovered between three large populations of interneurons in the mouse visual cortex: 1. Interneurons expressing parvalbumin, which strongly inhibit each other but weakly inhibit other populations. 2. Interneurons expressing somatostatin, which do not inhibit each other but strongly inhibit all other neuron types. 3. Interneurons expressing vasoactive intestinal peptide, which preferentially inhibit somatostatin interneurons. Furthermore, Bergoin et al. (2023) examined various types of interactions between excitatory and inhibitory neurons and concluded that inhibitory neurons act as “stabilizers” in flexible neural networks, ensuring the long-term retention of information acquired during learning.

However, working with combinations of such neurons, we concluded that combinations of these three types of neurons (excitatory, inhibitory, and “prohibition”) are insufficient to model certain important operations, such as the continuous movement of activity patterns through neural networks. However, such operations are clearly quite common during rapid processing in the visual brain. For example, they are essential for the functioning of visual working memory and for rapid visual recognition.

To overcome this problem, we hypothesized that some kind of synchronization of processing occurs in the neural modules of the visual brain. Specifically, we hypothesized that each neural module contains an ordered impulse activity in the form of a kind of “clock frequency” controlled by “endogenous pacemakers” (EP).

Having carried out a computational experiment with our proposed three-layer neural network with synchronization, we found that in this case the preservation of the pattern shape is ensured by slowing down the blurring at the outer and inner boundaries of the pattern.

Figure 4 shows a specific version of the neural network we synthesized above – a neural module for recording the activity pattern received for processing in order to prevent it from being “blurred.”

**Fig. 4.**
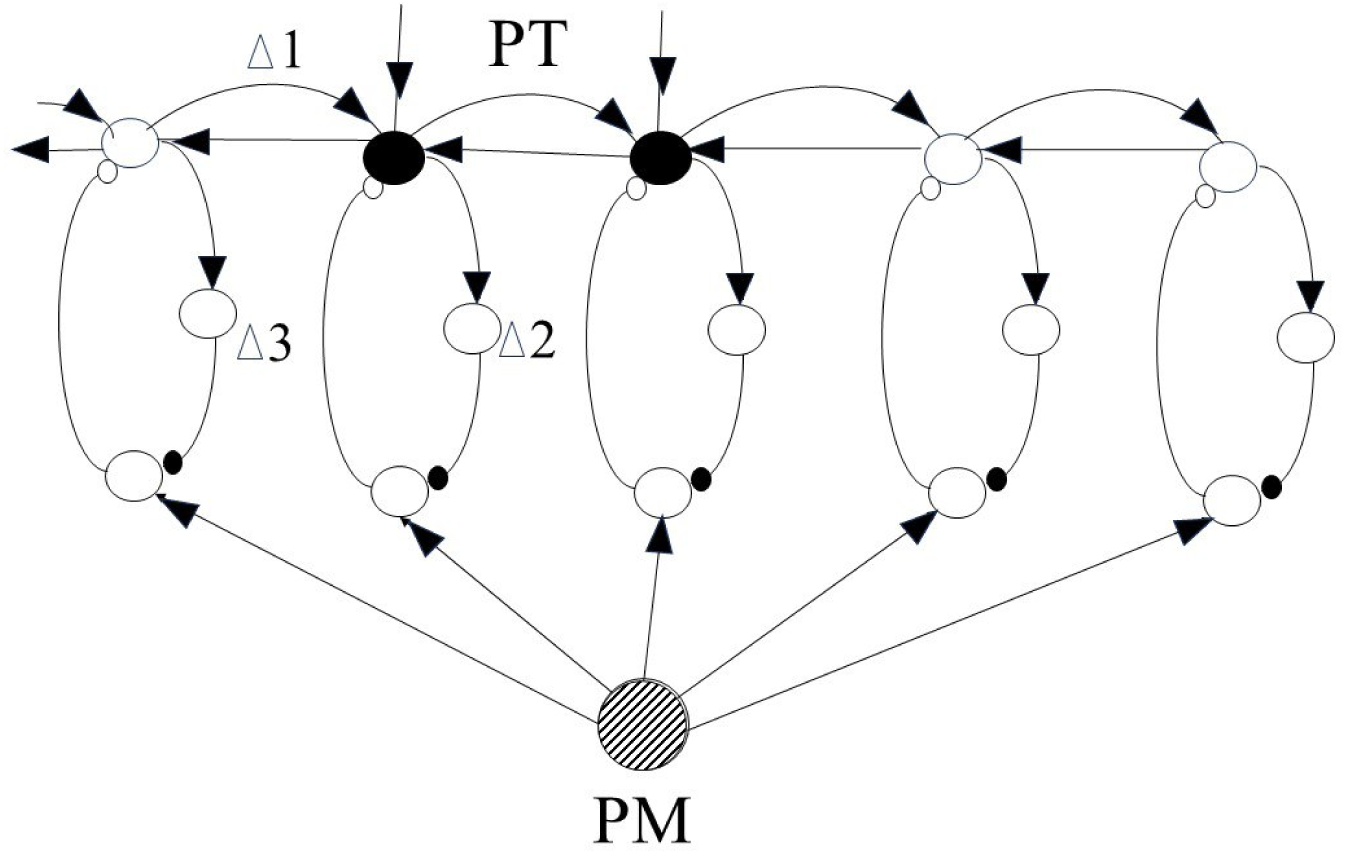
A neural module for capturing the pattern of activity in the visual cortex corresponding to the input image on the retina. Only a fragment of this extended three-layer neural network is shown. This circuit uses two types of inhibitory neurons: 1. regular ones, which act on excitatory neurons; 2. special “prohibition” neurons, which interact only with inhibitory neurons of the first type, locking (disabling) them at certain points during pattern processing. The PM is a pacemaker that operates at the gamma rhythm frequency, and the pattern corresponding to the input image is fed to the input at the same frequency. The subtlety of this entire network lies in the fact that the phase shifts of all these rhythms must be correctly selected (i.e., the delays Δ1, Δ2, and Δ3 must be correctly set). In our experiment, the following condition was met: Δ1>Δ3>Δ2.

The operating principle of this network is as follows. In the “fixation” mode, inhibitory neurons in the second layer inhibit neurons closest to the edges of the pattern, preventing it from losing its shape and spreading left and right. The pattern remains fixed because excitatory connections from its neurons turn off corresponding inhibitory neurons in the second layer of the module (“below the pattern”). Modeling using the Deepsik AI showed that this network reliably fixes the pattern and protects it from blurring (see the repository https://github.com/victoriasafiulina-design/ModelV).

### Modeling a neural module for pattern translation during visual recognition, in working memory, and in the hippocampus

A classic requirement for visual recognition is pattern invariance under affine transformations (shifts, similarities, and rotations), i.e., maintaining its topological connectivity (Levashov, 2021). Figure 1 shows the pattern invariance. Figure 5 shows a slightly modified neural network compared to the one shown in Figure 4, in order to use the added synchronization bus to move the pattern left or right in accordance with commands “from above”.

**Fig. 5.**
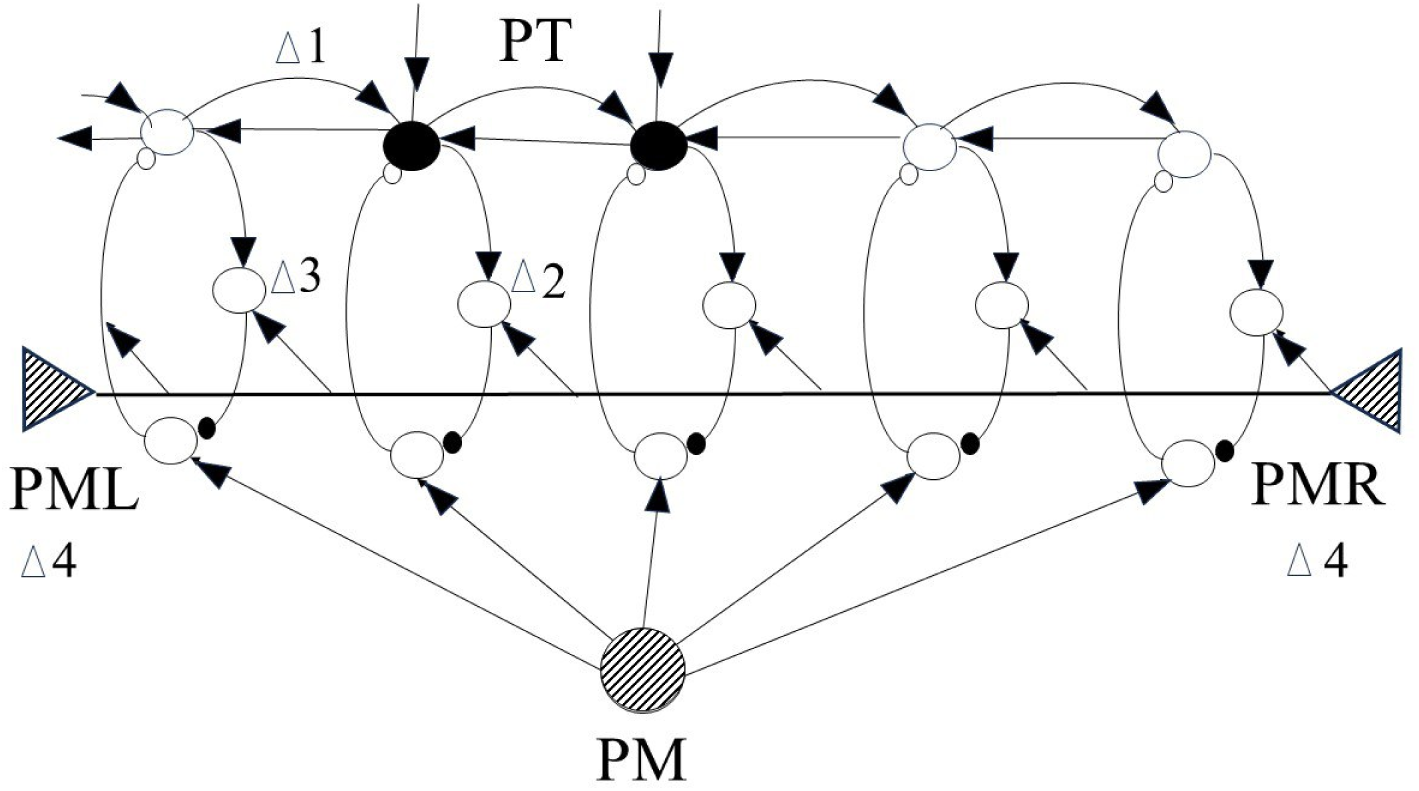
Schematic diagram of the neural module for shifting a pattern to the right or left. Only a fragment of the extended three-layer neural network is shown (the one-dimensional case or side view, as in Fig. 4). In this diagram, as before, two types of inhibitory neurons are used: 1. regular ones, which act on excitatory neurons, and 2. “prohibition” neurons, which interact only with inhibitory neurons of the first type, locking (disabling) them at certain moments during pattern processing. The original pattern (filled circles) can be shifted to the right along the neural network by synchronizing with the gamma rhythm frequency coming from the right along the bus from the PMR pacemaker (shown in the figure on the right). To shift to the left, the second pacemaker, PML (shown in the figure on the left), must be activated. When activated, both pacemakers operate at the gamma rhythm frequency (100 Hz), and the pattern corresponding to the input image is fed to the input at the same frequency. The ratio of delays (phase shift of gamma rhythm oscillations) is also important here: Δ4 > Δ1> Δ3> Δ2.

We conventionally define a “bus” as a serial connection of identical neurons in a single cortical layer within a single neural functional module. In this scheme, rhythmic activity from one of two pacemakers is transmitted along the “bus.” Rightward pattern shift requires synchronization from the PMR, while leftward pattern shift requires synchronization from the PML.

A key moment in our modeling was an experiment in which a “bus” for gamma synchronization was added to the neural module. This made it possible to implement a controlled linear shift of the pattern while maintaining its topological integrity. The modeling results demonstrate that synchronization of this specialized neural module with the help of endogenous pacemakers enables the execution of operations on a discrete neural ensemble as a holistic object. These results suggest that this may be the mechanism by which the visual cortex operates during recognition, visual working memory operations, and the hippocampus.

Modeling using the Deepsik AI showed that this network allows for leftward or rightward shifts of the activity pattern (see Appendix).

Is our proposed “clock synchronization” mechanism physiological? We believe that the result obtained in the work of Bhattacharya et al. (2022) may provide evidence for such a mechanism. In this work, it was shown that activity oscillations in the prefrontal cortex are organized as traveling waves over a wide frequency range, and that they propagate in a linear fashion.

### Hypothesis on neural functional modules and their synchronization during pattern processing in the visual system

Above, we demonstrated that a neural module with two types of inhibitory neurons enables the processing of topologically related, but essentially discrete, patterns of activity in the visual brain. From a technical standpoint, each such operation must be performed strictly synchronously and very quickly, almost simultaneously with all elements of the pattern. This is because all events during visual processing in the brain last a fraction of a second, with patterns on the retina changing every 300-400 ms. This is a necessary condition, since the processed pattern itself is simply a discrete set of excited, albeit interconnected, neurons.

In this article, we propose to consider the visual brain as a hierarchical structure of neural functional modules with the presence of synchronization in each of them.

#### 1. Neural functional modules (NFM)

We believe that each operation in visual recognition is performed by individual NFMs. Such a module is a local neural ensemble performing only one or two operations. Individual NFMs in the visual brain are combined into larger functional units, for example, a visual form recognition unit, a motion perception unit, or a stereo perception unit. The necessity of such a hierarchical structure is dictated by the organism’s need to survive in our extremely complex, dynamic visual environment (Levashov, 2018, 2021).

This hypothesis has a number of physiological confirmations. For example, in a study by Malvache et al. (2016), they examined the CA1 hippocampal region in actively moving mice. Using two-dimensional calcium imaging, they identified patterns of synchronous neuronal activity that indicate the presence of functionally independent (orthogonal) neural ensembles. Here, the authors use the term “orthogonal neural ensembles” in a mathematical sense. This means that the identified neural ensembles are functionally independent and weakly correlated with each other in activity. In other words, each such ensemble implements a distinct pattern of co-activation of neurons. Each time the animals performed different tasks, they activated, demonstrating discrete temporal segments of activity.

In another article, van Es et al. (2025), analyzing magnetoencephalography data from healthy humans, the authors found that the activity of large functional brain networks is organized into stable cyclic patterns. Several states of the cerebral cortex were identified, each representing a unique spatial configuration of activity. These states correspond in location to known functional blocks of the cortex—the visual perception block, the passive mode block, the dorsal block of spatial attention, etc. The authors concluded that key cognitive functions such as attention, memory, visual perception, and others are not activated randomly, but rather occur in a strictly ordered cycle lasting 300-1000 ms.

Further evidence supporting the presence of localized NMFs in the cortex comes from modeling data from Antonello et al. (2022). In this study, they analyzed bursts of spikes recorded from cultured individual hippocampal neurons in rat embryos aged 6 to 35 days. They examined how the network topology changes during maturation. They found that the number of connections in the networks increased, demonstrating structural integration. According to the authors, the networks developed by forming modules, with most modules including neighboring neurons (as in the hypothetical neural modules we proposed above).

#### 2. The Role of Synchronization in Visual Brain Operations

The most obvious example of the significant role rhythmic activity plays in pattern processing is the behavior of the alpha rhythm in the occipital cortex. It exhibits a distinct 8-10 Hz rhythm with the eyes closed, but disappears in most subjects upon opening the eyes. From an engineer’s perspective, this suggests that the alpha rhythm can only be detected “idling,” when the system is in the “standby” state. When the work process is activated, its synchronicity disappears, since the different units of any machine or technical system each operate at their own pace and are activated only at the right time. The idea that the alpha rhythm acts as a kind of “clock frequency” for the visual system was first proposed by Norbert Wiener, the father of cybernetics.

Thus, in this article, we describe hypothetical neural modules whose modeling revealed that synchronization enables the solution of a number of tasks useful for the visual brain.

In the visual brain, different operations require different processing speeds. For example, the sensorimotor response to suddenly appearing stimuli, such as the appearance of a large moving object, should be processed by neural modules with the shortest processing cycles. Operations involved in visual recognition of objects, especially when the observer is moving, should also have a short cycle. Conversely, some working memory operations, such as those associated with low reading proficiency, may be performed at a lower clock rate. Overall, the range of processing cycle durations in the visual cortex can be estimated as ranging from 100-150 ms (a rapid visual or visuomotor response to a significant stimulus) to 500-1000 ms (working memory operations, decision making in the presence of alternatives). Interestingly, these figures coincide with the range of duty cycles of the functional blocks described in the above-mentioned work of van Es et al. (2025).

## General Discussion

We will discuss a key point in our work: the role of rhythm synchronization in processing activity patterns in living sensory networks.

### Possible mechanisms for generating endogenous pacemaker rhythms

We propose that synchronization of processing in each neural module can be ensured by “endogenous rhythms” (ER). As discussed above, such rhythms can be considered a kind of analog to the clock frequency in electronic devices.

We will assume that each neural module has its own pacemaker mechanism that generates the required “clock frequency.” Different neural modules can process input stimuli either in parallel or sequentially, depending on the commands received “from above” (presumably from the frontal cortex). In this case, we can draw an analogy between our neural modules with their clock frequencies and the motor centers that generate various cyclical movement patterns—walking, running, and dance movements (Hooper, 2000). However, analogies alone are not enough. It’s necessary to understand how, from a physiological perspective, ER generation might occur. In other words, how is the clock frequency generated for each neural module?

We suggest that the most likely mechanism is the “ephaptic” generation mechanism. Since the neuron membrane is a dielectric, rhythmic changes in the external electric field (for example, the alpha or theta rhythm) will cause oscillations in the membrane potential. If a neuron is close to the action potential threshold, such rapid changes in membrane potential can “push” it to fire. Such a neuron becomes a kind of “slave pacemaker”—it generates discharges in phase with the external rhythm, for example, in phase with one of the external electrical rhythms recorded in EEG, or with the “endogenous” weak fields of neighboring firing neurons.

Indeed, there are in vitro studies on brain slices or neuronal cultures in which, when weak electric field oscillations are applied, neurons synchronize their activity with the field oscillation rhythm. In one of the first studies of this type, Jefferys et al. (2003) used guinea pig hippocampal slices exposed to an alternating electric field of 1-5 V/m. If the neurons in the slice were sufficiently densely packed, the changes in the transmembrane potential of these neurons were almost proportional to the distance from the radiation source. The authors found a direct proportionality between the number of excited neurons and the field strength.

Later, the same group of authors, in a study performed on rat hippocampal slices, demonstrated that a low-amplitude alternating electric field (1-5 mV/mm) can modulate the spiking activity of neurons, synchronizing it in phase with the applied field (Deans et al., 2007). A similar conclusion was reached in the work of Fröhlich and McCormick (2010), where the authors showed that the effect of even weaker alternating electric fields (0.2-1 mV/mm) on neocortical slices can also evoke and synchronize the active states of the neural network with the frequency of the applied field.

Finally, Anastassiou et al. (2011) showed that weak extracellular electric fields (∼1 mV/mm), mimicking the fields generated by neural tissue itself, can modulate spiking activity and synchronize the firing of individual neurons and neural ensembles in cortical slices, providing direct evidence for an ephaptic mechanism of such modulation.

Currently, we cannot say for sure which frequencies are used in neural modules as endogenous rhythms. However, it is quite possible that their frequencies coincide with the electrical rhythms known in physiology and recorded as EEG. In this case, endogenous rhythms may well be “imposed” by external EEG rhythms. The opposite is also possible—that the activity of neighboring neurons that arises during development gradually synchronizes and can be collectively recorded as an EEG rhythm.

### Spontaneous Emergence of Rhythms in Neural Network Models

It turns out that there is evidence of spontaneous synchronization occurring early in development. For example, Sun et al. (2010) used electrode arrays to record the activity of individual neocortical neurons in mice aged 6 to 15 days in vitro. These experiments showed that the number of neurons with spiking activity increased significantly between 6 and 15 days after birth, and by day 9, synchronized network activity emerged, which later became the primary discharge pattern. These results indicate that individual neurons can self-organize into complex neural networks for information processing early in development.

One of the pioneers in the study of brain rhythms is D. Buzsáki. According to his theory, the gamma rhythm in the brain’s neural networks is formed through the interaction of excitatory pyramidal neurons and inhibitory interneurons, when conditions arise for “fast inhibition” of the perisomatic regions of these neurons via GABA receptors (Buzsáki and Wang, 2012). According to Buzsáki’s idea, during the narrow, 3-ms window of the gamma cycle, excited neurons have time to turn off inhibitory neurons. When inhibition of this local network ceases, a new round of excitatory neuron firing occurs. Thus, there is only a short window for excitation (3 ms), while inhibition dominates during the longer phase of the gamma cycle.

In an earlier study by Buzsáki and Wang (1996), the authors performed a computer experiment on over a hundred neurons with spiking activity. Modeling showed that the presence of inhibitory interneurons in such networks allows for the generation of gamma rhythms, but only in the presence of a constant tonic external excitatory input.

However, later, the authors of another modeling study, Tikidji-Hamburyan et al. (2015), demonstrated that the system of coupled oscillators implemented in the Wang-Buzsáki model is extremely sensitive to inhomogeneity of the excitatory input. If neurons in the network received slightly different excitatory inputs, synchronization was easily disrupted. This contradicted previously obtained experimental data, as precisely this type of inhomogeneity is observed in neurophysiological studies.

Another modeling study is the article by Traub et al. (2000). In this study, a network of 3,072 pyramidal cell analogues and 384 interneurons was modeled. The goal was to determine the conditions under which oscillations similar to those observed in neurophysiological studies could arise in this network. This model explained the generation of oscillations by the presence of GABA receptors and axon-axon electrical contacts. The model also predicted that action potentials during sustained gamma rhythms originate specifically in the axons of pyramidal cells.

## Conclusion

In the visual brain, different operations require different processing speeds. For example, the visuomotor response to suddenly appearing stimuli, such as a large moving object, should be processed by neural modules with the shortest processing cycles. Operations involving visual recognition of stationary objects and simultaneous observer movement also require short cycles. Levashov (2018), recording subjects’ initial eye movements to images with ambiguous meanings, demonstrated that cortical processing of such complex objects lasts in cycles of only 50-80 ms. After such processing, subcortical structures accurately direct the gaze to the most informative next point of interest. The literature contains numerous articles demonstrating how different rhythms correlate with the performance of various cognitive tasks. For example, the gamma rhythm is more pronounced and synchronized during the processing of sensory information and the control of motor acts. In contrast, lower-frequency rhythms (alpha and beta, 8-30 Hz) are characteristic of tasks requiring executive control and monitoring by the frontal cortex (Fries, 2005, 2015; Miller et al., 2018).

In this article, we emphasized that the local neural structures that perform all the necessary pattern manipulations in visual recognition and other similar processes are controlled by endogenous synchronizing rhythms. This raises the question: what are the rhythms recorded on the EEG? Are they a net derivative of endogenous rhythms, or, conversely, are endogenous rhythms “inductively induced” by the primary EEG rhythms? At this point, it is impossible to say definitively which is primary and which is secondary. It is just as difficult to answer the question “What comes first—the chicken or the egg?”

## Summary

In this article, we used a so-called “neuroengineering approach” to studying the functioning of living sensory systems, based on the synthesis of neural structures with continuous logic (Pozin, 1971; Pozin et al., 1978) and their computational modeling (Levashov, 2021). We described and modeled the synchronization mechanism of a neural network based on endogenous pacemakers, which, similar to electronic devices, generate clock frequencies for processing input stimuli. We demonstrated that, in the presence of “endogenous synchronizing pacemakers” in a neural module, continuous operations on discrete sets of active neurons can be performed in the visual brain.

We consider the results obtained here important for two reasons:

1. They suggest that this may be the neurobiological mechanism underlying recognition, visual working memory, and hippocampal function. These neural structures are associated with the rapid processing of extended, complex visual patterns.
2. At the same time, our approach is also relevant to artificial intelligence, particularly the development of intelligent autonomous robots. Such developments currently use computer vision and deep neural network-based training, which is quite resource-intensive and does not provide movement and recognition accuracy comparable to that of living prototypes. Our neural modeling may open up new possibilities for building neuromorphic intelligent architectures as an alternative to costly deep learning neural networks.

## Data availability

All relevant data generated or analyzed during this study are included in this published article. Numerical source data for generating the figures are available. The custom neural network simulation code (ModelV) used for the experiments is accessible in a public GitHub repository: https://github.com/victoriasafiulina-design/ModelV.

## Article and author information

### Author details

**OLEG LEVASHOV**

Institute of Control Sciences, Russian Academy of Science, Moscow, USSR, neuroengineer Research Center of Neurology, Russian Academy of Science, Moscow, Russia, researcher Brain & Body Development Centre, Da Lat City, Lam Dong Province, Vietnam, Leading researcher

**Contribution:** Conceptualization, Methodology,Formal analysis,Validation, Writing, Visualization

**Competing interests:** No competing interests declared

**VICTORIA SAFIUINA**

Institute of Physiology of Komi Science Centre of the Ural Branch of the Russian Academy of Sciences (IPhys FRC Komi SC UB RAS)

**Contribution:** Modeling,Investigation, Software, Validation, Formal analysis, Writing,Visualization

**Competing interests:** No competing interests declared

## Application

**Fig. A.**
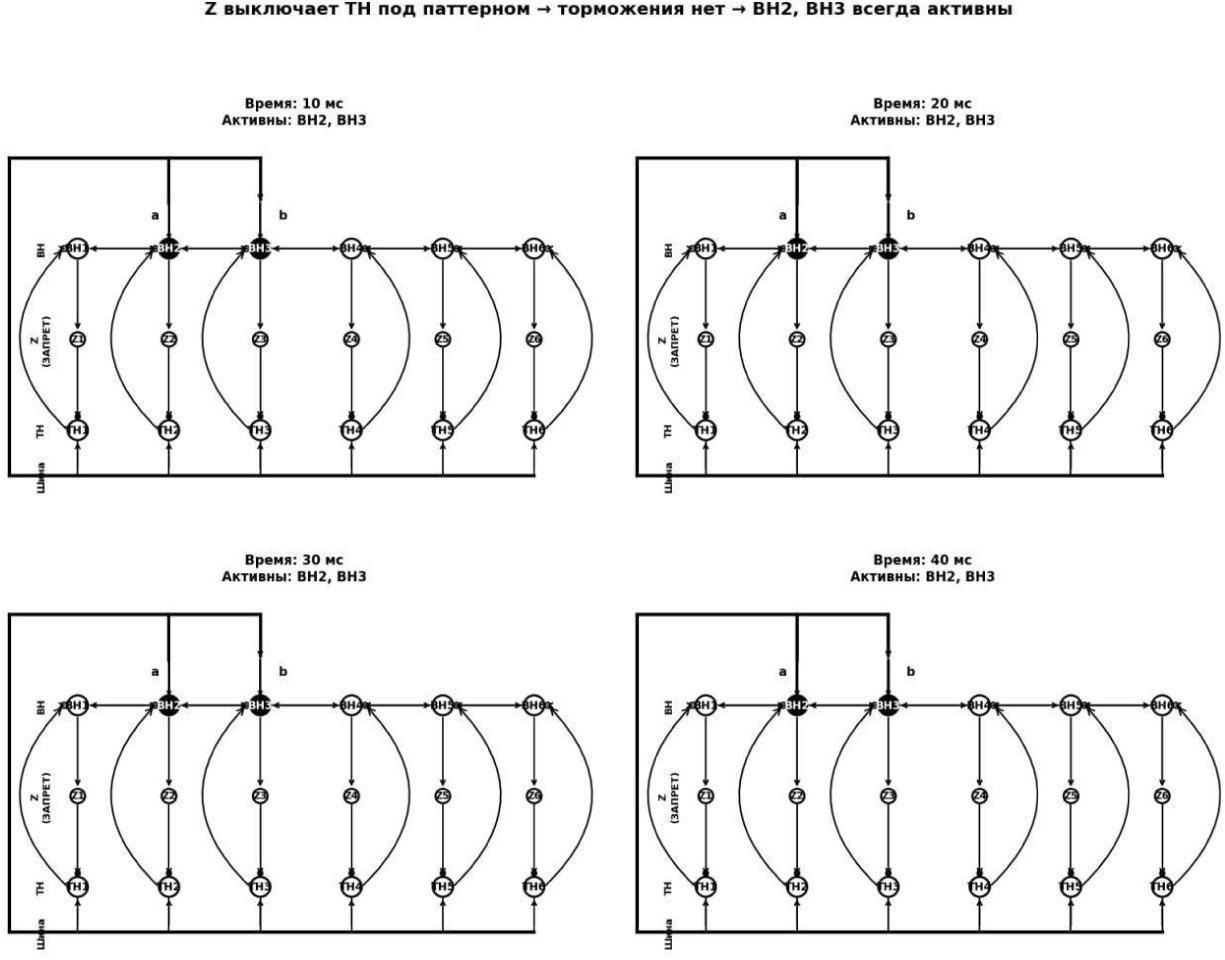
NS for fixing (freezing) a pattern.

**Fig. B.**
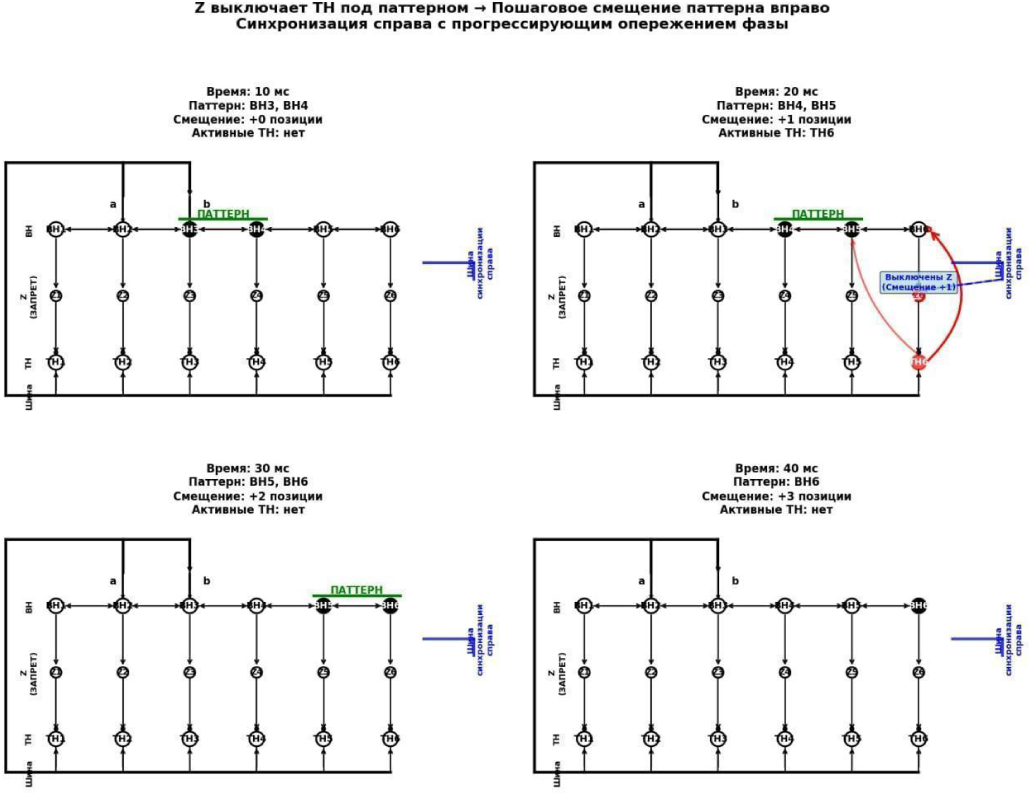
NN for moving a pattern along a neural layer.

